# A specialized bacterial group II intron is a highly efficient retrotransposon

**DOI:** 10.1101/2024.12.23.630110

**Authors:** Lucie Gomes, Claire Toffano-Nioche, Daniel Gautheret, Maria Costa

## Abstract

Mobile group II introns are site-specific retrotransposons composed of a large self-splicing ribozyme and an intron-encoded reverse transcriptase that are widespread in bacterial and organellar genomes. Sequence and structural variations of the ribozyme and the associated reverse transcriptase define several lineages of bacterial group II introns. Interestingly, some of these intron families evolved different mobility strategies while others colonize particular genetic contexts. Here, we have investigated the mobility activity of an *Escherichia coli* group II intron that is inserted into the stop codon of the stress-response gene *groEL*. Using mobility assays based on over-expression from a donor plasmid, we demonstrate that this intron is a highly efficient and site-specific retrotransposon, capable of colonizing the *groEL* gene of an *E. coli* host strain according to the insertion pattern observed in natural genomes. Furthermore, we provide evidence that a chromosomal copy of the full-length retrotransposon can be expressed from its native genetic locus to yield mobile retroelement particles. This intron constitutes a novel model system that could help reveal original mobility strategies used by some group II intron retrotransposons to colonize bacterial genomes.

## INTRODUCTION

In bacteria, mobile group II introns constitute the most abundant class of retrotransposons. These composite mobile elements result from the association of a large self-splicing catalytic RNA (ribozyme) with a multi-functional reverse transcriptase (RT) enzyme encoded by the intron itself (Lambowitz and Zimmerly, 2011). The group II intron ribozyme folds into a characteristic and evolutionarily conserved secondary structure organized into six domains (I to VI). Phylogenetic analyses of the intron RNA structures define six major structural subtypes (IIA1, IIA2, IIB1, IIB2, IIB, IIC) (Michel et al., 1989; Toor et al., 2001) which, as a general rule, co-evolve with specific phylogenetic lineages of the encoded reverse transcriptase (Toor et al., 2001; Zimmerly et al., 2001; Simon et al., 2008).

Group II intron retrotransposons do not carry their own promoter, therefore, they are transcribed from any adjacent host promoter as part of the precursor RNA transcript. Then, in association with its RT enzyme, the ribozyme core catalyses self-splicing of the entire intron in the form of a large lariat RNA carrying a typical 2’-5’ branch structure (Saldanha et al., 1999; Costa et al., 2016; Haack et al., 2019). The canonical mobility pathway of these retroelements, known as ‘retrohoming’, is initiated by reverse splicing of the excised lariat into double-stranded DNA targets (Zimmerly et al., 1995a; Cousineau et al., 1998) follow by cDNA synthesis by the intron-encoded reverse transcriptase according to the ‘target DNA-primed reverse transcription’ mechanism (TPRT; (Zimmerly et al., 1995b)). The vast majority of bacterial group II intron lineages recognize their DNA targets with a very high degree of sequence specificity, based primarily on the multiple Watson-Crick base pairs that intron RNA establishes with the top strand of the DNA target (Moran et al., 1995; Eskes et al., 1997; Mohr et al., 2000). The intron RNA sequences engaged in these base pairings lie in domain I and are called ‘EBS’ (Exon Binding Sites) while the complementary DNA target sequences are known as ‘IBS’ (Intron Binding Sites). Because the intron EBS sequences involved in mobility are also used for base pairing with segments located in the flanking 5’ and 3’ exons during self-splicing (Jacquier and Michel, 1987), group II intron retrotransposons invade DNA targets having ‘ligated-exons’ sequences. The RT component of the mobile RNP contributes to target recognition through stabilization of specific RNA-DNA interactions (Monachello et al., 2021) and supposedly, by making direct, yet-unidentified, contacts with a few other positions of the DNA target (Singh and Lambowitz, 2001; Jiménez-Zurdo et al., 2003; Zhuang et al., 2009a; Mohr et al., 2010).

Analyses of the distribution of group II introns in bacterial genomes reveal that the vast majority of introns are inserted into intergenic regions of the genome or inside other mobile genetic elements such as IS, transposons, integrative or conjugative plasmids (Dai and Zimmerly, 2002; Novikova et al., 2014; Waldern et al., 2020). This insertional pattern, typical of selfish mobile DNA, ensures survival of the element through minimal interference with host functions and provides opportunities for propagation within and between genomes. Interestingly, a few intron subgroups have evolved particular structural features that allow them to target specific intergenic contexts. This is the case of the well-studied subgroup IIC introns that recognize DNA hairpin structures and form short EBS-IBS pairings in order to insert downstream rho-independent transcriptional terminators (Granlund et al., 2001; Robart et al., 2007; Mohr et al., 2018). Nevertheless, and in stark contrast with the insertional pattern described above, a few bacterial introns are found inserted into crucial housekeeping genes of the host and, at least in some reported cases, their splicing is involved in modulating expression of the host gene (Chee and Takami, 2005; Simon et al., 2008; McNeil and Zimmerly, 2014; Pfreundt and Hess, 2015). In an instructive example, in the cyanobacterium *Trichodesmium erythraeum*, splicing of the four closely related ORF-less group II introns interrupting the *dnaN* gene (encoding the β subunit of DNA polymerase III) is under the control of a structurally related ORF-carrying intron present in the *RIR* gene (encoding the ribonucleotide reductase) (Meng et al., 2005). The RT expressed from the full-length *RIR* intron functions as a general maturase, assisting splicing of all the intron variants. Therefore, in this example, group II intron splicing results in coordinated expression of the *RIR* and the *dnaN* genes (Meng et al., 2005). Another striking example of bacterial group II introns specifically associated with protein-coding genes is provided by a monophyletic family of subgroup IIB1 introns whose members are found inserted within, or in immediate vicinity to, initiation or termination codons of housekeeping genes (Michel et al., 2007). All members of this lineage also carry a large 5’-subterminal insertion (∼300 - 700 nt) located within the small internal loop at the base of domain I. With the exception of their location in the secondary structure, these large insertions share no significant sequence similarity, except between the most closely related introns (Michel et al., 2007). Both the potential function of this extra domain and the process by which these peculiar introns colonize the start or stop codons of housekeeping genes are currently unknown. The only representative of this specialized clade that has been studied experimentally so far is the *Azotobacter vinelandii* IIB1 intron, Avi.groEL, which is inserted between the second and third position of the termination codon of the essential heat-shock *groEL* gene (encoding GroEL, an Hsp60 chaperonin) (Adamidi et al., 2003; Ferat et al., 2003). This integration pattern, which is conserved among all the relatives of the Avi.groEL intron, results in the replacement of the stop codon (UAA) of the *groEL* gene by the alternative stop codon (UAG) upon intron invasion (Michel et al., 2007). Previous biochemical work has shown that the Avi.groEL intron self-splices *in vitro* in the absence of its encoded reverse transcriptase (Adamidi et al., 2003; Ferat et al., 2003), however, its mobility potential has never been investigated. In this work, we have carried out the first characterization of the retrohoming activity of a novel ‘groEL’ intron variant found in the genome of *Escherichia coli* strain KTE56.

## RESULTS

### The specialized subgroup IIB1 intron EcKTE56.groEL is a highly active retrotransposon

The novel EcKTE56.groEL intron, which is inserted between the second and third position of the stop codon of the *groEL* gene in *Escherichia coli* strain KTE56 was identified using BLASTn to search the NCBI ‘wgs’ and ‘refseq_genomes’ databases for relatives of the previously reported *Azotobacter vinelandii* or *Citrobacter rodentium* groEL introns (Michel et al., 2007) in *Escherichia coli* genomes. The queries used for database mining consisted of the last 62 nucleotides of the *groEL* gene followed by the complete sequence of each groEL intron. These bioinformatics searches reveal that variants of the groEL intron are widespread among gammaproteobacteria (detailed results will be reported elsewhere).

The secondary structure of the EcKTE56.groEL ribozyme (Figure 1) is typical of bacterial subgroup IIB1 introns and harbors canonical forms of the tertiary interactions characteristic of this subgroup, including the recently demonstrated EBS2a-IBS2a base-pairing interaction between a site in the intron ‘EBS2’ loop and position -14 of the 5’-exon (Monachello et al., 2021). Moreover, and as all the other members of the IIB1 specialized clade, the EcKTE56.groEL intron carries a large 5’-subterminal insertion (hereafter designated as ‘subdomain Ia’) located within the internal loop at the base of domain I (Figure 1). In addition to its ribozyme moiety, the EcKTE56.groEL intron codes for a CL1-lineage RT containing a C-terminal endonuclease domain. Interestingly, our analyses reveal that, except for the subdomain Ia, the primary sequence of the EcKTE56.groEL intron is very similar to that of EcI5, a well-characterized *Escherichia coli* subgroup IIB1 intron (Zhuang et al., 2009a; Monachello et al., 2021). Nevertheless, the EcI5 intron lacks any insertion at the base of domain I and does not belong to the monophyletic family associated with initiation or termination codons. The RT enzymes encoded by the EcKTE56.groEL and the EcI5 introns share ∼56% of amino acid identity and their ribozyme moieties have ∼70% sequence identity in pairwise alignment (the alignment was performed on RNA intron sequences deleted from their domain IV section; in addition, the entire subdomain Ia was further deleted from the EcKTE56.groEL sequence).

**Figure 1.**
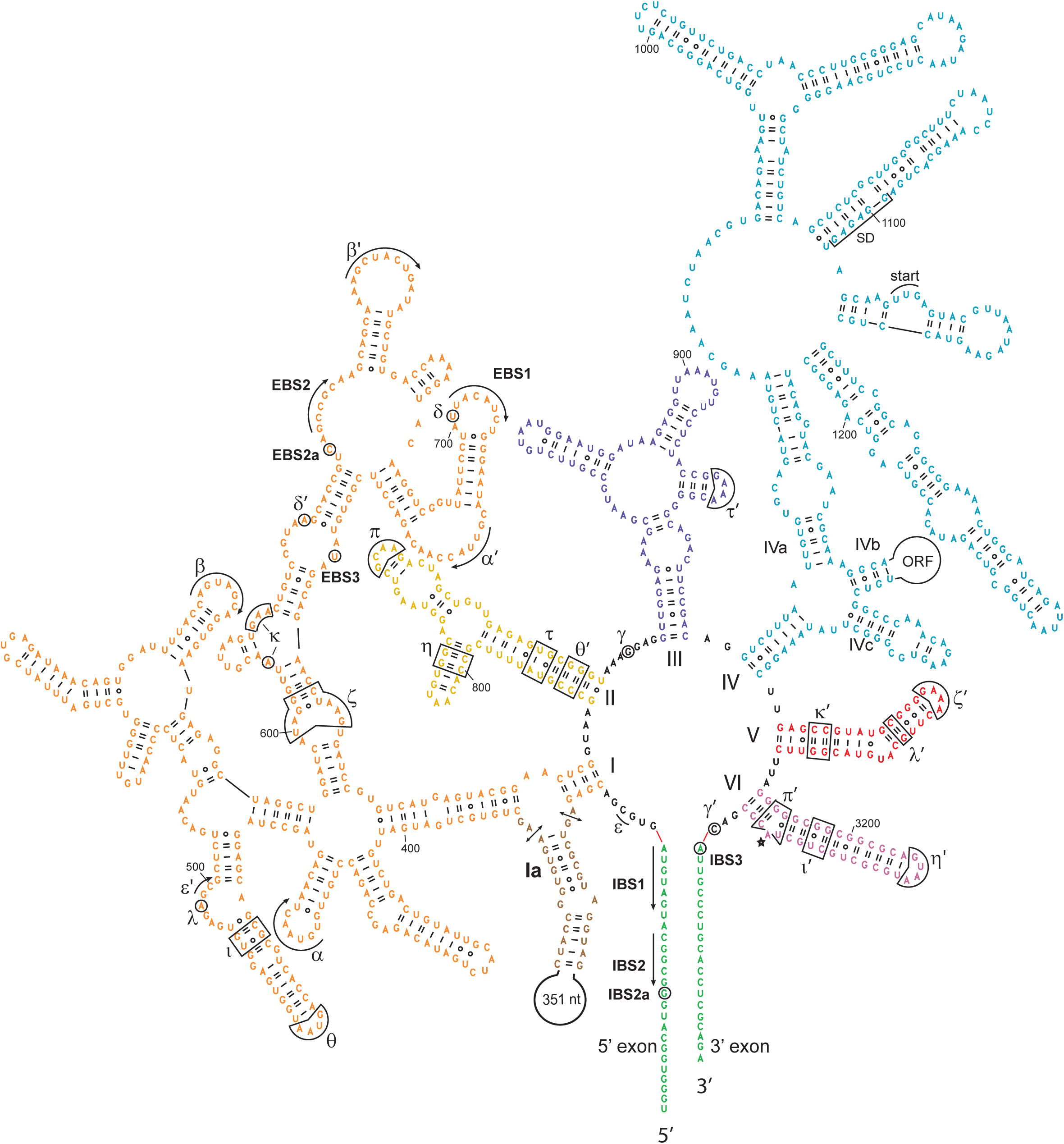
Secondary structure of the *Escherichia coli* subgroup IIB1 intron EcKTE56.groEL. The secondary structure of the EcKTE56.groEL intron is colored by major domains (I to VI). Tertiary base-pairing interactions between the intron and its 5′ and 3′ flanking exons are designated as EBS (Exon Binding Site) - IBS (Intron Binding Site) pairings whereas long-range interactions involving intron sequences are designated by greek letters. Subdomain Ia is colored in brown and the double arrows mark the ends of the segment deleted in mutant construct EcKTE56.groEL_ΔIa (see Materials and Methods). The IVa subdomain contains the high affinity binding site for the EcKTE56.groEL RT enzyme and is present in the ribozyme construct EcKTE56.groEL_ΔORF that is mobilized by the intron RT expressed in tandem from the same donor plasmid ‘pACD4K_EcKTE56.groEL-IntronΔORF + RT’ (see Materials and Methods). Subdomain IVa also contains the translation signals used for synthesis of the RT: the Shine-Dalgarno sequence (SD) and the UUG start codon (start). Red lines connecting the intron and flanking exon sequences represent the splice sites.

In order to investigate the retrohoming activity of the EcKTE56.groEL intron, we first adapted our previously described *Escherichia coli*-based mobility assay used for the EcI5 intron (Monachello et al., 2021). This assay is based on a Cam^R^ donor plasmid that allows expression of the intron flanked by its 5’ and 3’ exons, from a T7lac promoter (Materials and Methods and Figure 2A). The donor plasmid is transformed into the *E. coli* host strain HMS174(DE3), which carries a chromosomal copy of the T7 RNA polymerase gene under the control of the IPTG-inducible promoter lacUV5. Importantly, this homologous expression system provides to the EcKTE56.groEL intron a perfectly matched genomic target site that corresponds to the end of the single-copy *groEL* gene of the host strain. This genetic context avoids the need to co-transform the cells with a recipient plasmid containing the appropriate intron target site. *E. coli* HMS174(DE3) cells transformed with the donor plasmid are grown in liquid culture in the presence of chloramphenicol and addition of IPTG to the culture induces over-expression of the intron, triggering mobility (Materials and Methods). We first investigated the mobility of the EcKTE56.groEL intron using two different intron configurations expressed from their respective donor plasmids (Materials and Methods). The first form corresponds to the native, full-length intron (EcKTE56.groEL_WT). In the second configuration, the intron is deleted of most of its ORF in domain IV but still carries subdomain IVa (corresponding to the beginning of the RT ORF), which contains the high-affinity binding site for the intron RT enzyme (Figure 1). This shorter intron version originates from a donor plasmid (‘pACD4K_EcKTE56.groEL-IntronΔORF + RT’) that expresses in tandem, both the EcKTE56.groEL_ΔORF intron flanked by its exons and the RT enzyme from a position downstream of the 3’ exon. The RT binds to the intron_ΔORF ribozyme and mobilizes it for genomic integration. In other group II intron systems, this much shorter version of the intron has been shown to increase very significantly the mobility frequency, probably because the smaller size of the intron_ΔORF RNA makes it less susceptible to degradation by the host nucleases (Guo et al., 2000). As a control for intron mobility through the canonical TPRT mechanism, we also tested a mutant version of the full-length intron encoding an inactive RT enzyme due to mutation of the YADD catalytic motif to YAHH (mutant EcKTE56.groEL_YAHH). In initial experiments, detection of retrohoming events occurring in bacterial populations was carried out by PCR on culture aliquots taken at different stages of the mobility assays (Materials and Methods). The agarose gel in Figure 2B indicates that retrohoming events occur at early stages of the mobility protocol, even before the addition of the IPTG inducer. Indeed, and despite the non-quantitative nature of the PCR reactions, the relative intensities of the DNA bands suggest that the full-length EcKTE56.groEL intron behaves as an efficient retroelement and that its mobility efficiency seems comparable to that of the smaller version of the intron EcKTE56.groEL_ΔORF (Figure 2B). In contrast, no PCR products were obtained for the EcKTE56.groEL_YAHH mutant, consistent with its inability to perform the ‘target DNA-primed reverse transcription’ step required for retrohoming. Sequencing of the PCR products corresponding to the 5’- and the 3’-integration junctions confirmed that both the full-length and the intron_ΔORF forms insert into the host genome between the second and third position of the stop codon of the *groEL* coding sequence (Figure 2C and Materials and Methods). This insertion pattern, which results in the change of the *groEL* termination codon from UAA to UAG, recapitulates the insertion pattern observed in gammaproteobacterial genomes for all the analysed groEL intron sequences (data not shown).

**Figure 2.**
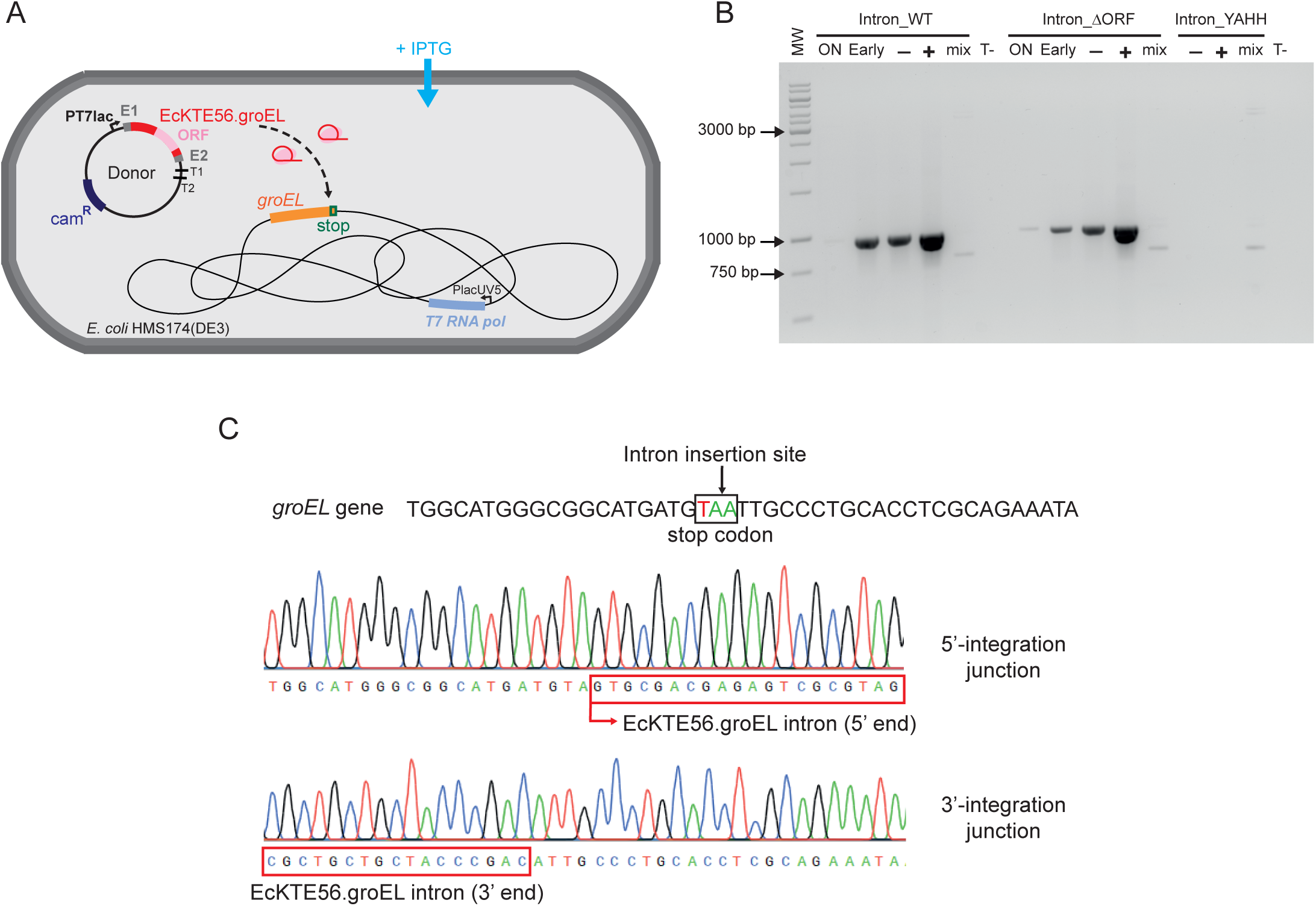
The EcKTE56.groEL intron is mobile in *E. coli*. (**A**) Diagram of the genetic system used for mobility experiments in *E. coli* host strain HMS174(DE3). Expression of the EcKTE56.groEL intron (wild-type or mutant versions) from the donor plasmid is triggered by addition of IPTG (Isopropyl β-D-1-thiogalactopyranoside) to the culture medium (for details, see text and Materials and Methods). (**B**) Representative agarose gel with PCR products indicating mobility events in bacterial cultures. *E. coli* HMS174(DE3) cells were transformed with the appropriate donor plasmid allowing expression of the specific intron construct indicated. The integration events into the chromosomal *groEL* gene were detected by PCR on culture aliquots taken at different stages of the mobility assay as follows: after overnight growth following transformation (‘ON’ lanes), after early log phase (‘Early’ lanes), after 90 minutes of incubation without IPTG (‘-’ lanes) or incubation with 100 µM of IPTG (‘+’ lanes). See also Materials and Methods for details. PCR amplification of the 5’-integration junctions was performed with primer ‘Sens ORFgroEL B’, which anneals to the terminal region of the host *groEL* gene, and intron-specific primer ‘Rev-3_KTE56Int1-1’ (primer sequences in Supplementary Table S1) and generates a 937-bp product. For each intron construct, the ‘mix’ lane is a control for PCR artifacts performed with the same primers as above and a mixture of 50 ng of *E. coli* HMS174(DE3) genomic DNA and 10 ng of each of the corresponding donor plasmids. The ‘T-’ lanes correspond to controls without culture lysate added to the PCR reaction. (**C**) Representative Sanger sequencing chromatograms of PCR products corresponding to the amplification of the 5’- and the 3’-integration junctions resulting from a mobility assay with the full-length EcKTE56.groEL_WT intron (the culture aliquot used for PCR was taken after 90 minutes of induction with 100 µM of IPTG). As indicated by the chromatograms, the intron inserts precisely between the second and third position of the stop codon of the host *groEL* gene.

In order to quantify the mobility efficiencies of the EcKTE56.groEL intron constructs, individual colonies plated at the end of the mobility assays were tested by PCR using primers that hybridize to the host *groEL* gene region, at sites flanking the intron integration site.

Figure 3A shows that under over-expression conditions (in the presence of 100 μM IPTG), the retrohoming frequency of the full-length WT intron is surprisingly high and comparable to that of the intron_ΔORF version. In contrast, no intron integration events were detected in similar assays performed with the TPRT-inactive EcKTE56.groEL_YAHH mutant, as expected. We also took advantage of the ‘leaky’ expression (i.e., basal expression of the gene under study that occurs in the absence of IPTG induction) characteristic of the phage T7 expression system on which our mobility assays are based, to test the above intron constructs in the absence of IPTG (Materials and Methods). Under these low-expression conditions, and as expected, the mobility efficiency of both intron constructs is significantly reduced. Nevertheless, the full-length WT intron again proves to be as mobile as the intron_ΔORF variant. Thus, our data demonstrate that the native EcKTE56.groEL intron is a highly active retroelement and that in contrast to previous reports for other mobile group II introns (Guo et al., 2000), the length of the intron RNA does not appear to be a primary determinant of EcKTE56.groEL mobility.

**Figure 3.**
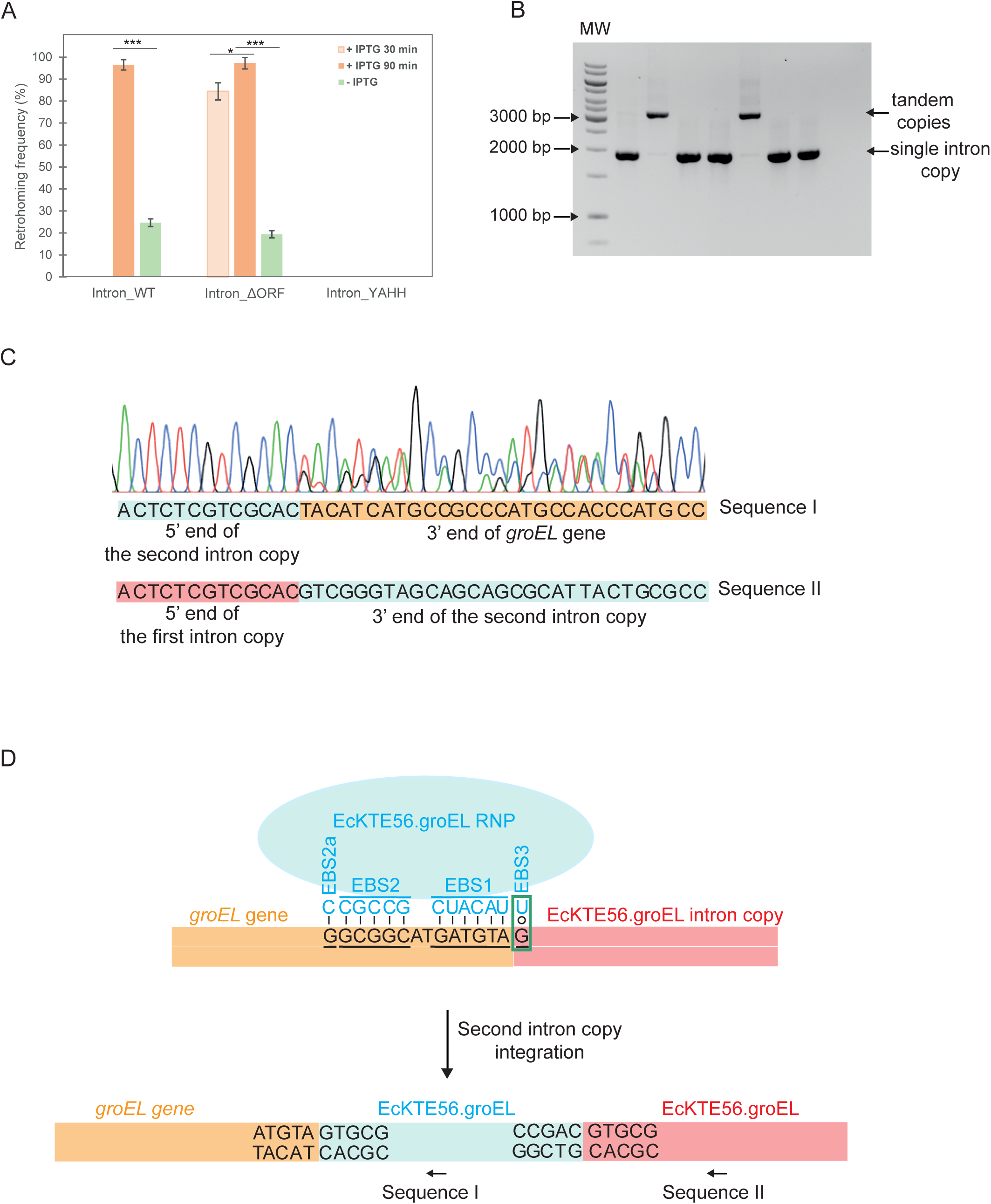
The full-length EcKTE56.groEL intron is highly mobile and the intron_ΔORF construct readily forms tandem intron copies at the chromosomal *groEL* locus. (**A**) *E. coli* HMS174(DE3) cells were transformed with the appropriate donor plasmid for expression of the specific EcKTE56.groEL intron construct indicated. Mobility assays were carried out under ‘over-expression’ conditions by addition of 100 µM IPTG for 30 minutes (light orange) or 90 minutes (orange) or under ‘leaky’ low-expression conditions with no IPTG added for 30 minutes (green; see Materials and Methods). After the mobility assays, cells were plated onto LB-agar + Cam + 20 mM glucose in order to obtain single colonies and retrohoming frequencies were calculated by colony PCR (Materials and Methods). The bar graphs show the mean of at least three independent experiments, with the error bars indicating the standard error of the mean. The stars represent two-sided P value with the unequal variances t-test (*<0.05, ***<0.001). (**B**) Representative agarose gel showing colony-PCR products corresponding to tandem integration of the EcKTE56.groEL_ΔORF intron into the host genome. Colony-PCRs were performed with forward primer ‘Sens ORFgroEL B’, which anneals to the terminal region of the host *groEL* gene, and ‘Rev K-12_avalGroEL’, which anneals downstream of the host *groEL* gene (primer sequences in Supplementary Table S1). Single-copy integration of the EcKTE56.groEL_ΔORF intron generates a 1914-bp PCR product whereas tandem integration of two intron copies generates a 3293-bp PCR product. (**C**) Representative Sanger sequencing chromatogram of the 3293-bp PCR products (tandem intron copies) shown in panel B obtained with reverse primer ‘Rev-3_KTE56Int1-1’ that allows to read the intron 5’-end. The chromatogram reveals the superimposition of two sequences (labelled ‘I’ and ‘II’) resulting from hybridization of the primer to the intron 5’-section in the two integrated copies (primer hybridization is represented by arrows in panel D). In sequences I and II, the segment corresponding to the terminal section of the host *groEL* gene is coloured in orange, the segment corresponding to the first intron copy integrated is in light red and the segments corresponding to the second intron copy integrated are in light blue. (**D**) Diagram of the EBS(RNA)-IBS(DNA) base-pairing interactions that mediate integration of the second intron copy. The EBS3(U)-IBS3(G) wobble pairing that involves the first nucleotide (G) of the first intron copy integrated is boxed in green. Elements are color-coded as in panel C. The sequence at the junction of the two intron copies (5’-..CCGACGTGCG..) is diagnostic of the tandem integration.

Interestingly, during screening of colonies for quantification of the retrohoming efficiency of the EcKTE56.groEL_ΔORF + RT system (in the presence of IPTG) some PCR reactions generated a product much larger than expected, whose estimated size was consistent with integration of two copies of the intron_ΔORF at the *groEL* locus in the host genome (Figure 3B). Sanger sequencing of this PCR product with a reverse primer that allows reading the intron 5’ end and its upstream context, shows the superimposition of two different sequences immediately upstream the intron 5’ end (Figure 3C). One of the two sequences corresponds to the end of the *groEL* gene while the other corresponds to the 3’ end of the intron. These results confirmed the tandem arrangement of the two integrated intron_ΔORF copies, as schematized in Figure 3D. It is important to note that this integration pattern is not a peculiarity of the intron_ΔORF construct since tandem copies were also detected for the full-length EcKTE56.groEL_WT construct, albeit at a much lower frequency (4.2% compared with 22.4% for the intron_ΔORF version; frequency among all colonies screened positive for the presence of at least one intron copy in the genome). The tandem integration pattern is most likely made possible by the composition of the EBS3 site of the EcKTE56.groEL intron, which, as for the vast majority of the groEL intron variants, happens to be a uridine. The presence of this U allows the formation of an EBS3(U)-IBS3(G) wobble pair involving the first intron nucleotide (G) installed at the end of the *groEL* gene by integration of the first intron copy. Although this U-G wobble pair is not the canonical EBS3-IBS3 interaction, it must support reverse splicing of the second intron copy to some extent, allowing intron insertion precisely at the junction between the end of the *groEL* gene and the beginning of the first integrated intron copy (Figure 3D).

A wealth of biochemical and genetic data has established that mobile group II introns are highly site-specific retrotransposons (Lambowitz and Zimmerly, 2011). Nevertheless, it was of interest to investigate the specificity of genomic integration of the EcKTE56.groEL intron under the experimental conditions used in our mobility assays. To that end, long-read nanopore technology was employed for whole-genome sequencing of genomic DNA prepared from overnight LB-cultures grown from a mixture of 73 independent colonies screened positive for the presence of a full-length intron copy in their genome. Analysis of all the reads that map to the host genome and cover, at least partially, the intron sequence revealed that all of them align exclusively to the end of the *groEL* gene and correspond to faithful integration events of the intron into the *groEL* stop codon (data not shown). Interestingly, a few reads (∼3% of all the intron-containing reads) were also seen to correspond to two full-length intron copies integrated in tandem, exactly as the pattern detected in our mobility assays described above. These results confirm that the EcKTE56.groEL intron moves exclusively by retrohoming, without generating ectopic integration events.

### Deletion of subdomain Ia significantly reduces intron mobility

The role of the large 5’-subterminal insertion, which is characteristic of the introns belonging to the specialized IIB1 clade, has remained unclear. Previous *in vitro* self-splicing experiments with the Avi.groEL intron ribozyme from *A. vinelandii* have shown that a construct deleted from its entire Ia subdomain self-splices significantly faster than the wild-type ribozyme mostly due to activation of the alternative hydrolytic pathway of splicing that yields linear intron (Ferat et al., 2003). We thus sought to determine whether the Ia subdomain of the EcKTE56.groEL intron could play a role in retrohoming. To that end, we use the previously described mobility assays to compare the retrohoming efficiencies of the EcKTE56.groEL_WT intron with that of the EcKTE56.groEL_ΔIa mutant, in which the entire subdomain Ia had been deleted (Materials and Methods). As shown in Figure 4, under IPTG-induction conditions, both retrotransposons exhibit similar levels of retrohoming. However, experiments carried out under low-expression conditions (absence of IPTG) reveal that the mobility of the EcKTE56.groEL_ΔIa mutant is in fact significantly lower, by about eightfold, than that of its wild-type counterpart. These results clearly indicate a role for the Ia subdomain at some stage of the retrohoming pathway that remains to be identified. Importantly, sequencing of the integration junctions for the EcKTE56.groEL_ΔIa retrotransposon showed that its specificity of integration into the groEL target site is not affected by the absence of subdomain Ia.

**Figure 4.**
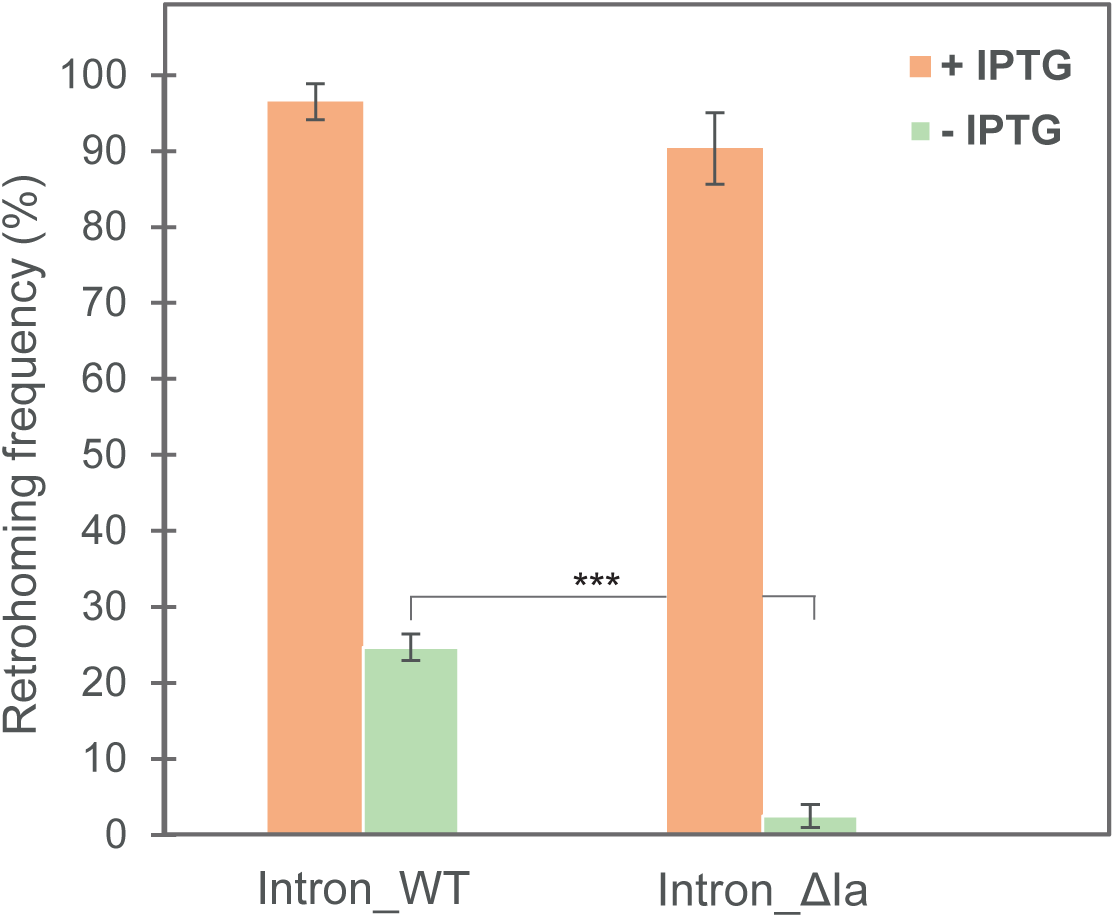
The EcKTE56.groEL_ΔIa mutant shows reduced mobility. *E. coli* HMS174(DE3) cells were transformed with the appropriate donor plasmid for expression of the specific EcKTE56.groEL intron construct indicated. Mobility assays were carried out under ‘over-expression’ conditions by addition of 100 µM IPTG for 90 minutes (orange) or under ‘leaky’ expression conditions with no IPTG added for 30 minutes (green; see Materials and Methods). After the mobility assays, cells were plated onto LB-agar + Cam + 20 mM glucose in order to obtain single colonies and retrohoming frequencies were calculated by colony-PCR (Materials and Methods). The bar graphs show the mean of at least three independent experiments, with the error bar indicating the standard error of the mean. The stars represent two-sided P value with the unequal variances t-test (***<0.001).

### An alternative over-expression system to study EcKTE56.groEL mobility in *E. coli*

In order to investigate the mobility of the EcKTE56.groEL retrotransposon in different *E. coli* strains without the need to install a copy of the phage T7 RNA polymerase gene in each host genome, we constructed a novel plasmid (pTara:500/pBR322; see Materials and Methods) containing the T7 RNA polymerase gene under the control of the arabinose-inducible promoter (pBAD). In addition, this novel plasmid carries an ampicillin-resistance gene and is compatible with the intron-expressing donor plasmid. Thus, after transformation of any desired *E. coli* strain with both the pTara:500/pBR322 and the intron-donor plasmid, over-expression of the retrotransposon can be induced by addition of arabinose to the culture (Figure 5A). Using this alternative system, we tested the mobility efficiency of the EcKTE56.groEL_WT intron in the MG1655 host, an *E. coli* K-12 laboratory model strain. The graph in Figure 5B shows that in this genetic context, the retrohoming efficiency achieved for the WT intron is as high as in the previous *E. coli* HMS174(DE3)/IPTG-inducible system which confirms the usefulness of our alternative arabinose-inducible system. Furthermore, and as in previous mobility assays, no integration events could be detected for the TPRT-inactive EcKTE56.groEL_YAHH mutant. The novel pTara:500/pBR322 plasmid may also constitute a good alternative to the commonly used pAR1219, a plasmid that also expressed the T7 RNA polymerase but from the IPTG-inducible lacUV5 promoter (Davanloo et al., 1984). Indeed, in our hands, liquid cultures of pAR1219-transformed cells grew consistently much slower than those of pTara:500/pBR322-transformed cells.

**Figure 5.**
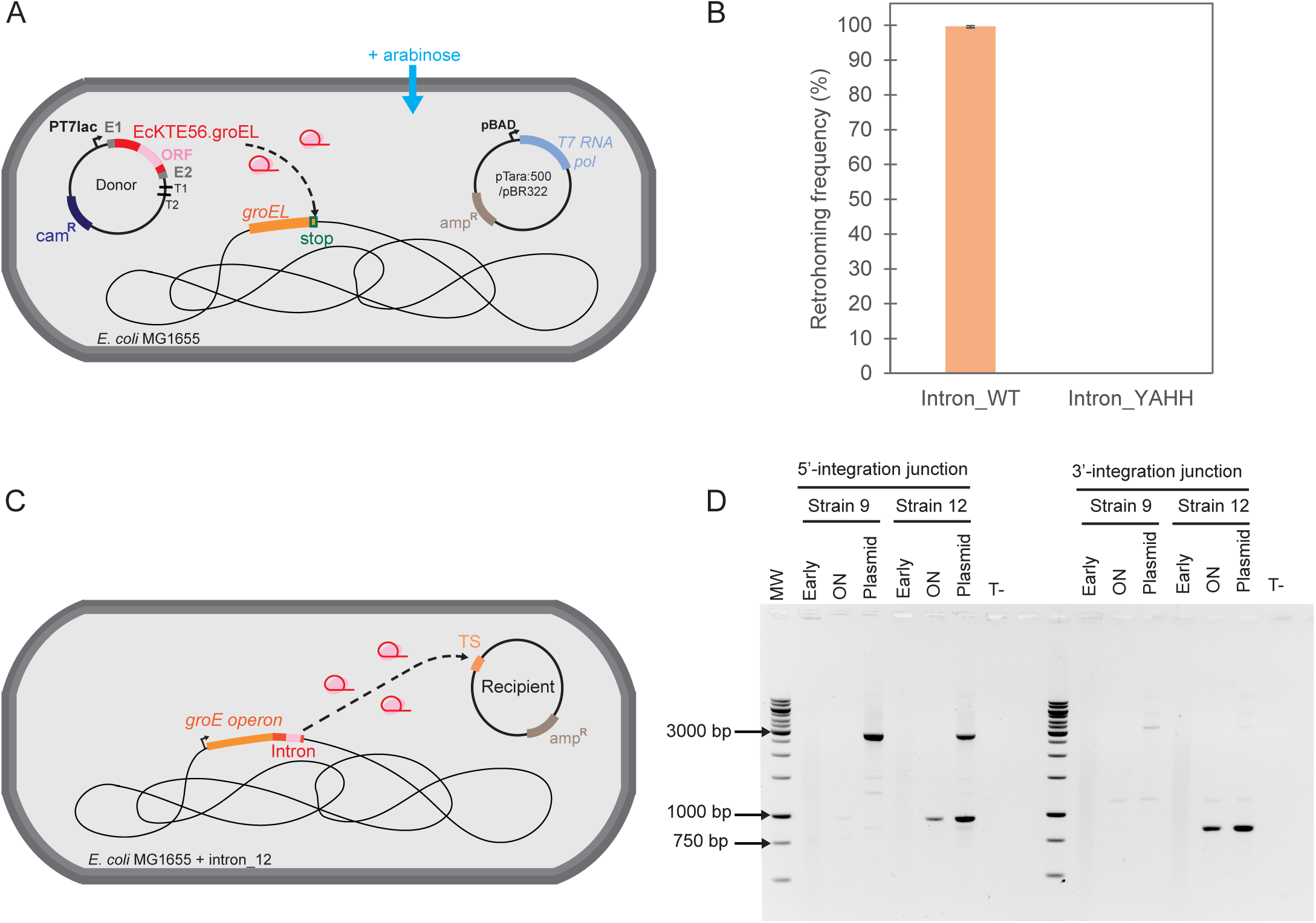
Mobility assays with another genetic system and from the groE operon. (**A**) Diagram of the alternative genetic system used for mobility experiments in *E. coli* MG1655. Addition of arabinose to the culture triggers transcription of the T7 RNA polymerase gene from the pBAD promoter carried by the novel plasmid pTara:500/pBR322, which then allows overexpression of the EcKTE56.groEL intron (for details, see text and Materials and Methods). (**B**) The pTara:500/pBR322 plasmid and the appropriate donor plasmid for expression of the specific EcKTE56.groEL intron construct indicated, were transformed into *E. coli* MG1655 by two successive electroporations. For mobility assays, cultures were induced by addition of 0.2% of L-arabinose (final concentration) for 90 minutes. After the mobility experiments, cells were plated on LB-agar + Cam + Amp + 20 mM glucose and retrohoming frequencies were calculated by colony-PCR (Materials and Methods). The bar graph shows the mean of at least three independent experiments, with the error bar indicating the standard error of the mean. (**C**) Diagram of the genetic system making use of a recipient plasmid with the target site (TS) for the EcKTE56.groEL intron that allows to detect potential intron mobility events generated from an intron copy integrated into the genome of *E. coli* MG1655 strain (only strain MG1655+Intron_12 is represented; see text and Materials and Methods). (**D**) Representative agarose gel with PCR products indicating intron mobility events into the recipient plasmid in the context of strain *E. coli* MG1655 + intron_12 (chromosomal wild-type intron copy) but not in the context of strain *E. coli* MG1655 + intron_9 (chromosomal mutant intron copy). Cells were grown at 37°C and PCR of the integration junctions were performed on culture aliquots taken at different stages as indicated: after early log phase, ‘Early’ lanes and after overnight growth, ‘ON’ lanes. The same type of PCR were also performed on purified recipient plasmid DNA extracted from the overnight cultures (‘Plasmid’ lanes; see Materials and Methods for details). The ‘T-’ lanes correspond to controls with no DNA added to the PCR reaction. PCR of the 5’-integration junctions were performed with forward primer ‘EcI5_13’, which anneals to the recipient plasmid backbone, and the intron-specific primer ‘Rev-3_KTE56Int1-1’ and generates a 953-bp product. PCR of the 3’-integration junctions were performed with reverse primer ‘EcI5_15’, which anneals to the recipient plasmid backbone, and the intron-specific primer ‘Sens dIV Eckte56Int1’ and generates a 850-bp product (primer sequences in Supplementary Table S1). The strong DNA bands around 3000 bp in the ‘Plasmid’ lanes of the 5’-integration junction PCRs were sequenced and found to be PCR artifacts due to nonspecific priming of the EcI5_13 primer only on the recipient plasmid backbone.

### Retrohoming activity generated from a chromosomal copy of the EcKTE56.groEL intron

Having demonstrated that the EcKTE56.groEL intron is a highly active retrotransposon when over-expressed from a donor plasmid it was of interest to try and determine whether a chromosomal copy of the intron could also be expressed from its native genetic location and generate a detectable retrohoming activity. In order to address this question, we first used the arabinose over-expression system described above to construct an *E. coli* MG1655 strain containing a single chromosomal copy of the EcKTE56.groEL_WT intron integrated at its natural locus. This new strain, hereafter named, ‘MG1655+Intron_12’ was then transformed with a recipient plasmid (pBR322_EcKTE56.groEL-TS_3XT2) that encodes the intron target site (Figure 5C and Materials and Methods). After transformation, liquid cultures were grown at 37°C and potential integration events into the recipient plasmid target site were monitored in the cell population by PCR with primers that allow amplifying either the 5’ or the 3’-integration junctions. In addition, PCR reactions were also performed on purified recipient plasmid DNA extracted from the overnight cultures. As a control, we performed the same experiments with strain ’MG1655+Intron_9’, which was fortuitously identified during whole-genome sequencing of candidate strains. In this strain the integrated intron copy contains three mutations, one of which is located in the RT domain of the enzyme and induces a premature stop codon that most probably prevents synthesis of a functional reverse transcriptase (Materials and Methods).

Consequently, this mutated copy should be unable to generate mobile RNP particles. The PCR results in Figure 5D show that mobility into the recipient plasmid occurs in the context of strain MG1655+Intron_12 since integration events are detected in the cell population as well as in the recipient plasmid DNA purified from the culture. In contrast, and as expected, there was no indication for mobility in the context of strain MG1655+Intron_9, which carries a mutated retrotransposon. Sequencing of the 5’ and 3’-integration junctions performed on the extracted recipient plasmid DNA confirmed the fidelity of the intron integration events. Although the retrohoming data collected in these experiments are not quantitative, they nevertheless clearly indicate that the wild-type EcKTE56.groEL retrotransposon can be expressed from its native locus. As the intron is not autonomous for its own transcription, its expression is certainly driven by the promoter sequences that govern transcription of the *groE* operon, which encompasses the *groES* and *groEL* genes. The *groE* operon is constitutively transcribed (at low-levels) at physiological temperatures and undergoes a transcriptional burst upon heat-shock (Roncarati and Scarlato, 2017). Since the above experiments were carried out at 37°C, our results show that the expression level of the chromosomal retrotransposon copy under physiological growth conditions is sufficient to induce significant retrohoming activity. When the above experiments were repeated with the addition of a heat-shock step at 45°C for 30 minutes, no significant differences were observed for PCR signals (data not shown).

## DISCUSSION

In this work we have carried out the first characterization of EcKTE56.groEL, a novel group IIB1 intron that colonizes the stop codon of the essential *groEL* gene in *Escherichia coli* strain KTE56. Our mobility assays, based on two different over-expression systems, demonstrated that the EcKTE56.groEL intron is a highly specific and active element, with a retrohoming frequency of ∼100% (under induction conditions) that ranks it among the most active introns studied to date, namely, the *Escherichia coli* EcI5 intron and the *Thermosynechococcus elongatus* Th.e.I3 intron, two other subgroup IIB1/CL1-lineage RT introns. It is remarkable that the EcKTE56.groEL intron achieves such high levels of mobility already in its native, complete form, despite its very large size (3228 nt). Indeed, RNA length does not appear to be a limiting factor for the mobility of this intron since shortening of the intron RNA (by deleting most of its ORF section) did not improve its efficiency, in contrast to what has been observed for other intron systems (Guo et al., 2000). In an attempt to rationalize our findings, it is worth recalling previous work demonstrating that ribosome particles (70S and 30S) can associate with the group II intron Ll.LtrB from *Lactococcus lactis*, both *in vivo* and *in vitro*, protecting the intron RNA from degradation by the *E. coli* 5′-endoribonuclease, RNase E (Contreras et al., 2013). It is tempting to propose that the high mobility of the EcKTE56.groEL intron may result, at least in part, from the same type of association with the ribosome, allowing the intron to efficiently escape RNA degradation.

Nevertheless, the association of the Ll.LtrB intron (belonging to subgroup IIA) with the ribosome has been shown to involve interactions with RNA structures in domains I and IV, most of which are not conserved in subgroup IIB1 introns (Contreras et al., 2013). Consequently, a potential association of the EcKTE56.groEL intron with the ribosome would have to rely on a rather different set of molecular interactions than those described for the Ll.LtrB intron.

Although it shares many sequence similarities with EcI5, the EcKTE56.groEL intron also has unique features, as it belongs to a specialized clade of group IIB1 introns that are specifically associated with start or stop codons of genes in proteobacteria. As all the other members of this family, the intron carries a large extra domain (subdomain Ia) inserted in the 5’ branch of the internal loop at the base of domain I (Figure 1). Our mobility assays under low-expression conditions revealed that subdomain Ia plays an important role in intron mobility, since its deletion leads to a decrease of about eightfold in retrohoming efficiency. Although the molecular mechanisms underlying this effect remain to be determined, several hypotheses can be put forward for the action of subdomain Ia. For example, subdomain Ia could induce some alternative intron RNA conformation(s) directly involved in a molecular mechanism of the retrohoming pathway specific to the EcKTE56.groEL intron (and possibly relevant to other groEL intron variants too). Nevertheless, the positive effect of subdomain Ia on retrohoming could also be indirect, via stimulation of the splicing process. Indeed, *in vitro* self-splicing results obtained for the *A. vinelandii* intron ribozyme Avi.groEL (also mentioned in the Results section) suggest that its Ia subdomain plays a role in promoting splicing via the branching pathway (lariat formation) at the expense of hydrolysis (Ferat et al., 2003). This effect on branching would have a favorable impact on retrohoming, since the lariat form is the only one capable of achieving complete and faithful reverse splicing (Zhuang et al., 2009b). Another hypothesis, which is not incompatible with the previous ones, is that subdomain Ia could constitute a specific binding site for one or more cellular factors (in line with the previous paragraph, the ribosome is one of the possible candidates). Such an association would probably not only offer some degree of protection against RNA degradation but could potentially participate in actively ‘guiding’ the intron RNP to its target site, i.e., the stop codon of the *groEL* gene, thus increasing retrohoming efficiency. A potential ‘guided-retrohoming’ mechanism would not be unprecedented, as the subgroup IIB intron RmInt1 from *Sinorhizobium meliloti*, is directed towards DNA replication forks (where it retrohomes very efficiently) via interaction with the β-sliding clamp factor, a component of DNA polymerase III (García-Rodríguez et al., 2019). In the case of RmInt1 however, the interaction with β-sliding clamp involves the intron RT enzyme and not the ribozyme component of the RNP. Although extensive genetic and biochemical studies are needed in the future to test all these hypotheses, the remarkable mobility activity of the EcKTE56.groEL intron makes it an excellent model system for effectively answering these important questions.

Another important result of our work is the finding that, once integrated into the stop codon of the host *groEL* gene, the EcKTE56.groEL intron remains an active retroelement able to produce RNP particles with retrohoming ability. Moreover, we found that intron expression happens already at physiological temperatures in the absence of any heat-shock stress. Although our results are qualitative, they clearly imply that the host *E. coli* RNA polymerase after transcribing the *groE* operon (*groES*-*groEL* genes) enters the intron sequence and continues synthesis to some extent, eventually generating some amount of full-length intron precursor transcripts that can then undergo processing through intron self-splicing. The occurrence of faithful retrohoming events into the recipient plasmid target site also implies that the EcKTE56.groEL intron self-splices from the *groESL*-intron precursor transcript through branching (at least in part), since as described above, the lariat form is the only one to operate the two transesterification reactions required for full-reverse splicing into the DNA target site (Zhuang et al., 2009b). Furthermore, it is interesting to consider that in the case where self-splicing and translation of the *groESL*-intron transcript occur in the same time window, ribosomes covering the *groEL* stop codon region (which houses the RNA structures of the 5’ end of the intron), will most likely affect intron RNA folding thus contributing to self-splicing regulation.

The ease with which retrohoming activity can be generated from the chromosomal copy of the EcKTE56.groEL intron in our laboratory *E. coli* strains, suggests that the intron in its natural genetic environment is also a highly active retrotransposon. This intron property, combined with the very high degree of sequence conservation of the *groEL* gene in proteobacteria (Kumar et al., 2015), suggests that the EcKTE56.groEL intron probably has a high potential for dissemination among natural gammaproteobacterial strains.

In conclusion, the novel EcKTE56.groEL intron constitutes a novel model system for future investigations aimed at discovering new mobility strategies evolved by group II introns to colonize specific genomic sites.

## MATERIALS AND METHODS

### Strains

*E. coli* strain DH5α was used for cloning. *E. coli* HMS174(DE3) strain (Novagen) was used for most of the intron-mobility experiments described in this work. This strain possesses the *recA1* mutation in a K-12 background and is suitable for the over-expression of desired target genes based on the phage T7 expression system. The ‘wild-type’ model *E.coli* K-12 strain MG1655 was used in some intron mobility experiments and to construct strains carrying a chromosomal copy of the EcKTE56.groEL retrotransposon as described below. All strains were grown in LB medium and when necessary, antibiotics were added to LB liquid medium or LB-agar at the following final concentrations: ampicillin (Amp), 100 μg/mL and chloramphenicol (Cam), 25 μg/mL.

### Plasmid constructs

The complete sequence of the wild-type EcKTE56.groEL intron flanked by 25 and 20 nucleotides of its natural 5’ and 3’ exons, respectively, was generated by gene synthesis (Eurogentec). For cloning purposes, in the synthesized gene, the EcKTE56.groEL precursor sequence was flanked, upstream by segment 5’-CTGCAGTGTACACAATTGAAGGACAACGCATATGGCTTCAG, and downstream by segment 5’-CCCGGGCATGCGGTACCTCGAGGATCC. In order to generate the ‘pACD4K_EcKTE56.groEL-IntronWT’ donor plasmid, expressing the complete wild-type EcKTE56.groEL intron and flanking exons, the synthesized DNA (provided inserted into vector pUC57) was digested with BsrGI and XhoI and ligated with plasmid pACD4K-C-loxP_Circ, previously linearized with the same restriction enzymes. The pACD4K-C-loxP_Circ plasmid, already described in Monachello et al. (2021), provides the backbone of the new donor plasmid containing a p15A origin of replication and a chloramphenicol resistance gene. The donor plasmids ‘pACD4K_EcKTE56.groEL-IntronYAHH’ and ‘pACD4K_EcKTE56.groEL-IntronΔIa’, expressing the mutant versions of the EcKTE56.groEL intron tested in this work were obtained from plasmid pACD4K_EcKTE56.groEL-IntronWT by cloning PCR products generated with the appropriate primers.

In order to build plasmid pACD4K_EcKTE56.groEL-IntronΔORF expressing the EcKTE56.groEL intron deleted of most of its ORF in domain IV, two PCR products were generated using plasmid pACD4K_EcKTE56.groEL-IntronWT as a template; one PCR (A) used primers KTE56DeltaORF5Fw and KTE56DeltaORF5Rv (primer sequences in Supplementary Table S1) and another PCR (B) was performed with primers KTE56DeltaORF3Fw and KTE56DeltaORF3Rv. PCR (A) was digested with AvrII and NsiI and PCR (B) was digested by NsiI and XmaI. After digestion these PCR products were ligated together with plasmid pACD4K_EcKTE56.groEL-IntronWT previously digested with AvrII and XmaI. In the final plasmid construct, only the first 127 nucleotides of the ORF were kept and the remaining ORF section was deleted (1867-nucleotide long deletion) and replaced by spacer 5’-ACGCGTAGTGTAATGCAT containing the MluI and NsiI sites.

The complete intron RT ORF with a 5’-adjacent phage T7 S10 Shine-Dalgarno sequence, was then cloned into plasmid pACD4K_EcKTE56.groEL-IntronΔORF downstream of the 3’-exon sequence. To perform this cloning step, we first generated a PCR product using primers KTE56-ORF_Fwd and KTE56-ORF_Rev and plasmid pACD4K_EcKTE56.groEL-IntronWT as the template. Then, this PCR product was introduced into plasmid pACD4K_EcKTE56.groEL-IntronΔORF with restriction enzymes SphI and XhoI to generate the final construct ‘pACD4K_EcKTE56.groEL-IntronΔORF + RT’.

The new ‘pTara:500/pBR322’ plasmid allows over-expression of the T7 RNA polymerase gene from the arabinose inducible pBAD promoter and carries the origin of replication and ampicillin resistance gene of vector pBR322. This new plasmid was obtained from combining pTara:500 (plasmid #60717 from Addgene), which carries the *araC* gene and expresses the T7 RNA polymerase from a pBAD promoter, with plasmid ‘pBR322_EcI5-TS_3XT2’, already described in Monachello et al. (2021), that has a pBR322 backbone. To generate the chimeric ‘pTara:500/pBR322’ construct, a PCR product was amplified from plasmid pTara:500 with primers pTara_araC_Fwd and pTara_T2_Rev while a second PCR product was amplified from plasmid pBR322_EcI5-TS_3XT2 with primers recEcI5_ampR_Rev and recEcI5_rop_Fwd. Then, both PCR products were digested with AatII and XhoI and ligated together.

The ‘pBR322_EcKTE56.groEL-TS_3XT2’ plasmid carrying the target site for the EcKTE56.groEL intron (the 5’ and 3’ sequences flanking the intron insertion site are 35- and 20-nt long, respectively) was constructed by the Gibson assembly method using plasmid pBR322_EcI5-TS_3XT2 as the backbone. This cloning step resulted in replacing the target site of the EcI5 intron present in plasmid pBR322_EcI5-TS_3XT2 with that for the EcKTE56.groEL intron.

All the constructs generated were verified by Sanger sequencing or by ONT whole-plasmid sequencing.

### Intron mobility experiments

For mobility experiments in *E. coli* HMS174(DE3), competent cells were transformed by heat-shock according to the manufacturer’s instructions, with 10 ng of each specific donor plasmid. After heat-shock, 80 μL of SOC media was added to the transformation reactions and the cells were incubated at 37°C for 1 hour to recover by growth. A small portion of this culture (10 μL) was serially diluted and plated onto LB-agar with chloramphenicol and 20 mM glucose (final concentration) in order to determine the efficiency of transformation. The remainder of the culture (90 μL) was added to 25 mL of LB medium containing chloramphenicol and 20 mM glucose (final concentration) followed by incubation at 37°C overnight. Glucose reduces the characteristic “leaky” expression (in the absence of IPTG) of the T7 phage expression system by catabolite repression on the lacUV5 promoter, which drives the T7 RNA polymerase gene. After overnight growth, the glucose was eliminated by washing the cells with 5 mL of fresh LB medium. Washed cells were resuspended in 15 mL of LB medium and a ∼100 μL portion of each washed culture was inoculated into 10 mL of LB medium containing chloramphenicol and incubated at 37°C until early log phase was reached (OD_590_ = 0.2 - 0.4). At this point, a 300 μL portion of each early log phase-culture was inoculated into glass culture tubes containing 3 mL of LB medium supplemented with 100 μM of IPTG (final concentration). Cultures were grown at 37°C for 90 minutes, or 30 minutes in some cases, as indicated in the figure legends. After this induction step, 1 mL of culture was used for OD_590_ measurement and the remaining 2 mL of culture was chilled on ice and then washed with 2 mL of ice-cold LB medium to eliminate IPTG (all centrifugations were performed at 4°C). Washed cells were resuspended in 2 mL of fresh LB medium, serially diluted and plated onto LB-agar + Cam + 20 mM glucose in order to obtain single colonies. Mobility experiments carried out under ‘leaky’ low-expression conditions were performed according to the same protocol as above except that after inoculation of a 300 μL portion of the early log phase-cultures into 3 mL of LB medium, no IPTG was added and the cultures were grown for 30 minutes before being serially diluted and plated onto LB-agar + Cam + 20 mM glucose in order to obtain single colonies.

Retrohoming frequencies were determined by colony-PCR with primers ‘Sens ORFgroEL B’ and ‘Rev K-12_avalGroEL’ (primer sequences in Supplementary Table S1) that hybridize to chromosome sequences flanking the intron integration site on the *groEL* gene of the *E. coli* HMS174(DE3) host. The size of the PCR products generated allow to discriminate ‘empty’ from ‘invaded’ *groEL* target sites and therefore, calculate mobility frequencies as the number of colonies having an integration event over the total number of colonies tested. For each intron construct, retrohoming frequencies were determined from at least three independent mobility experiments with (at least) a total of 90 colonies tested by PCR.

For detection of retrohoming events at the bacterial population level, PCR reactions were performed on lysates of culture aliquots (1 mL) taken at the end of the different steps of the mobility protocol as indicated in figure legends. The lysates were prepared by pelleting the bacterial cells, followed by resuspension in 200 μL of water and lysis at 95°C for 10 minutes. Then, cellular debris were pelleted by centrifugation at 15300 rpm for 2 minutes and the lysate supernatants were recovered. Each of these ‘diagnostic’ PCR probed approximately 5 x 10^6^ bacterial cells (calculated from cultures’ OD using relationship OD_600_ of 1.0 = 8 x 10^8^ cells/mL). PCR of the 5’-integration junctions were carried out using forward primer ‘Sens ORFgroEL B’, which anneals to the terminal region of the host *groEL* gene, and intron-specific primer ‘Rev-3_KTE56Int1-1’. PCR of the 3’-integration junctions were performed with primer ‘Rev K-12_avalGroEL’, that anneals downstream of the host *groEL* gene, and intron-specific primer ‘EcKTE56Int1_Fwd-1’ (for all the intron constructs except the intron_ΔORF construct). In order to amplify the 3’-integration junction for the intron_ΔORF construct, primer ‘Rev K-12_avalGroEL’ was used with the intron-specific primer ‘KTE56_D-IVa_Fwd’. The resulting PCR products were purified and sequenced in order to verify the 5’- and the 3’-integration junctions.

For mobility experiments in *E.coli* K-12 strain MG1655, cells were first electroporated according to (Figueroa-Bossi et al., 2022) with 10 ng of the new plasmid ‘pTara:500/pBR322’ that over-expresses the T7 RNA polymerase gene from the arabinose-inducible pBAD promoter, and transformants were selected on LB-agar + Amp. From these, competent cells were prepared for a second electroporation with the appropriate *cam^R^*-donor plasmid expressing the desired intron construct. Bacterial transformants containing both plasmids were selectively grown on LB-medium + Amp + Cam + 20 mM glucose (overnight, 37°C) and then, used for mobility experiments carried out under the same conditions as those described above, with the exception that, in this latter system, intron over-expression was triggered by addition of 0.2% of L-arabinose (final concentration).

### Construction of an *E.coli* MG1655 strain with a chromosomal copy of the EcKTE56.groEL retrotransposon

A new strain, named ‘MG1655+Intron_12’, was obtained from a typical mobility experiment based on the arabinose-induction system we developed, with *E.coli* MG1655 host strain transformed with plasmid pTara:500/pBR322 and donor plasmid pACD4K_EcKTE56.groEL-IntronWT by two successive electroporations, as described above. After the mobility experiment, cells were plated on LB-agar + Cam + Amp + 20 mM glucose. Several colonies were then streaked on LB-agar + Cam + Amp + 20 mM glucose and tested by PCR with primers ‘Sens ORFgroEL B’ and ‘Rev K-12_avalGroEL’ to identify clones containing a copy of the intron integrated into the stop codon of the host *groEL* gene. Curation of both plasmids was achieved by repeatedly growing the cells in LB medium with 20 mM glucose (typically, a 3 mL-culture was started by inoculating 3 μL of the previous culture and grew for 8 to 16 hours; about six successive cultures were grown over three days). Then, cells were serially diluted and plated onto LB-agar. To check for loss of both plasmids, cells from LB-agar plates were streaked on LB-agar + Cam and LB-agar + Amp plates. The genome of a few candidate strains was entirely sequenced by the nanopore ONT sequencing technology in order to verify the integrity of the retrotransposon sequence and the absence of mutations elsewhere on the chromosome. Except for the presence of the retrotransposon copy, strain MG1655+Intron_12 has no additional mutations compared with the parental MG1655 strain. Sequencing of another strain, named ‘MG1655+Intron_9’, revealed that the integrated retrotransposon copy contains three different mutations. A G-to-T mutation is found in domain IVa (upstream the ORF section) at intron position 1071. A second mutation corresponds to the insertion of four As at position 2049 in the ORF section that encodes the RT domain, closely downstream of the catalytic YADD motif. This insertion causes a frameshift that induces a premature stop codon 28 nt downstream of the frameshift, which should prevent the synthesis of a functional RT enzyme. The third mutation (T-to-C) is located downstream the frameshift mutation, in the ORF section that encodes the RT maturase domain.

### Intron mobility from a chromosomal copy of the EcKTE56.groEL retrotransposon

The new MG1655+Intron_12 strain was transformed by electroporation with 10 ng of the recipient plasmid pBR322_EcKTE56.groEL-TS_3XT2 according to (Figueroa-Bossi et al., 2022). As a control, strain MG1655+Intron_9 carrying a mutated retrotransposon copy, was also electroporated with the same recipient plasmid and underwent the same experimental protocol as strain MG1655+Intron_12. After electroporation, 1 mL of LB medium was added to the transformation reactions and the cells were incubated at 37°C for 90 minutes to recover by growth. Then, a small portion of the culture (10 μL) was diluted and plated onto LB-agar + Amp in order to determine the efficiency of transformation. The rest of the culture (990 μL) was centrifuged and concentrated into 100 μL and then added to 30 mL of LB medium + Amp followed by incubation at 37°C until early log phase was reached (OD_590_ = 0.2 - 0.4). At this point, a small portion of this culture (100 μL) was added to 20 mL of LB medium + Amp followed by incubation at 37°C overnight. Aliquots (4 - 6 mL) of overnight cultures were used for recipient plasmid extraction using the NucleoSpin plasmid kit (Macherey-Nagel).

Integration events of the EcKTE56.groEL retrotransposon into the recipient plasmid target site were detected by PCR on lysates of culture aliquots collected after the early log phase (10 mL) or overnight growth (3 mL). The lysates were prepared by pelleting the bacterial cells, followed by resuspension in 50 μL of water and lysis at 95°C for 10 minutes. Then, cellular debris were pelleted by centrifugation at 15300 rpm for 2 minutes and the lysate supernatants were recovered. Each one of these ‘diagnostic’ PCRs probed approximately 5 x 10^8^ bacterial cells (calculated from cultures’ OD using relationship OD_600_ of 1.0 = 8 x 10^8^ cells/mL). The recipient plasmid DNA extracted from the overnight cultures at the end of the experiment was also analysed by PCR for intron integration. In this case, 54.6 ng of purified plasmid DNA was used in each PCR reaction. Amplifications by PCR of the 5’-integration junctions were carried out with primer ‘EcI5_13’ (which anneals to the recipient plasmid backbone) and the intron-specific primer ‘Rev-3_KTE56Int1-1’. PCR of the 3’-integration junctions were performed with primer ‘EcI5_15’ (which anneals to the recipient plasmid backbone) and the intron-specific primer ‘Sens dIV Eckte56Int1’. The PCR products amplified from the extracted recipient plasmid DNA were purified and sequenced in order to read the 5’- and the 3’-integration junctions.

## ACKNOWLEDGEMENTS

M.C. acknowledges funding from program ‘Living machines at work’ (Université Paris-Saclay). We are indebted to Benoit Madec for his help with processing of ONT sequencing data.

---

**Supplementary Table S1.**
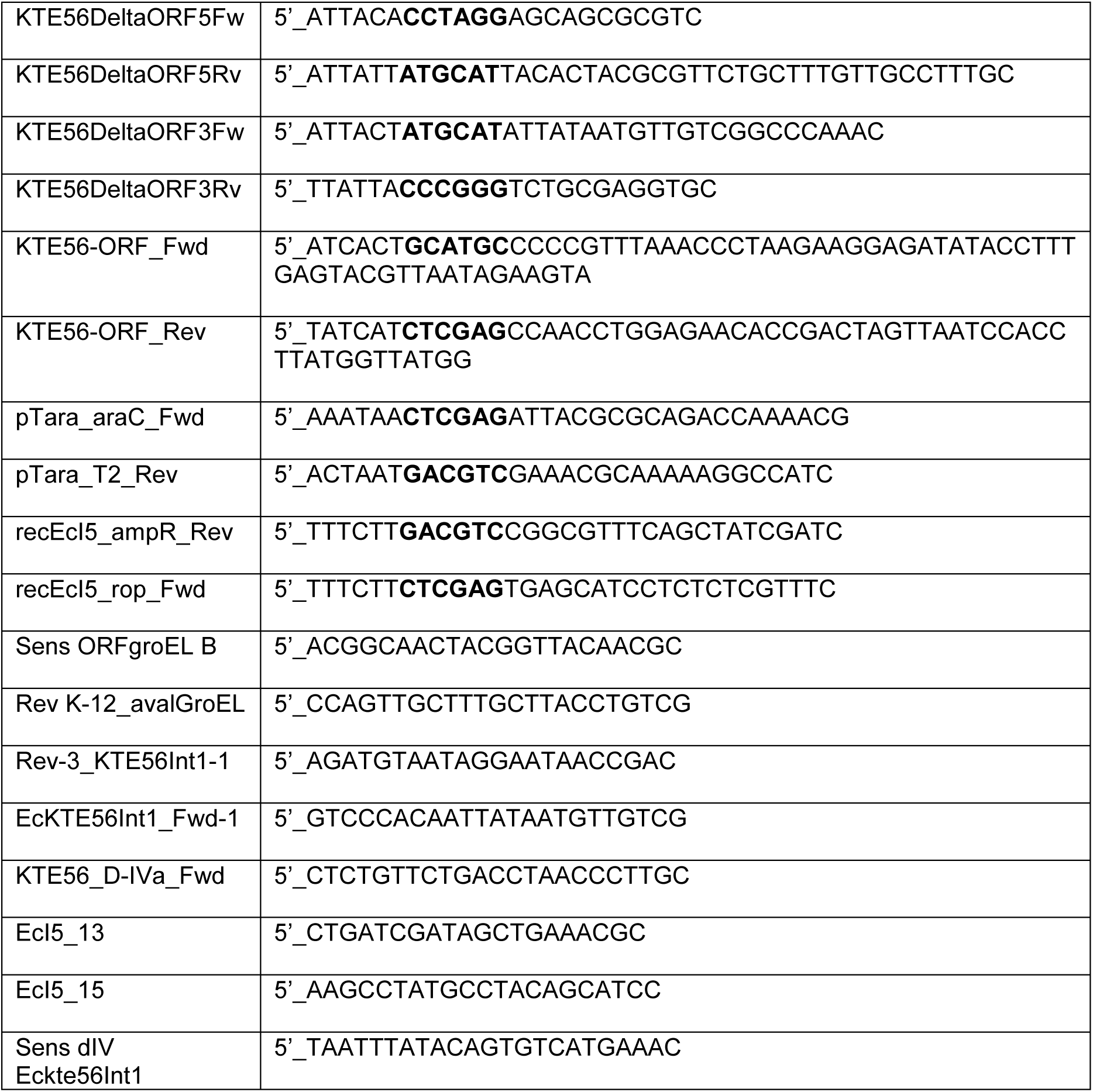
DNA oligonucleotides used for constructing the plasmids (sequences in bold indicate the restriction sites used for cloning) or for performing retrohoming-detection PCR as described in Materials and Methods.

## REFERENCES

Adamidi, C., Fedorova, O., Pyle, A.M., 2003. A group II intron inserted into a bacterial heat-shock operon shows autocatalytic activity and unusual thermostability. Biochemistry 42, 3409–3418. 10.1021/bi027330b

Chee, G.-J., Takami, H., 2005. Housekeeping recA gene interrupted by group II intron in the thermophilic Geobacillus kaustophilus. Gene 363, 211–220. 10.1016/j.gene.2005.08.017

Contreras, L.M., Huang, T., Piazza, C.L., Smith, D., Qu, G., Gelderman, G., Potratz, J.P., Russell, R., Belfort, M., 2013. Group II intron-ribosome association protects intron RNA from degradation. RNA 19, 1497–1509. 10.1261/rna.039073.113

Costa, M., Walbott, H., Monachello, D., Westhof, E., Michel, F., 2016. Crystal structures of a group II intron lariat primed for reverse splicing. Science 354, aaf9258. 10.1126/science.aaf9258

Cousineau, B., Smith, D., Lawrence-Cavanagh, S., Mueller, J.E., Yang, J., Mills, D., Manias, D., Dunny, G., Lambowitz, A.M., Belfort, M., 1998. Retrohoming of a bacterial group II intron: mobility via complete reverse splicing, independent of homologous DNA recombination. Cell 94, 451–462. 10.1016/s0092-8674(00)81586-x

Dai, L., Zimmerly, S., 2002. Compilation and analysis of group II intron insertions in bacterial genomes: evidence for retroelement behavior. Nucleic Acids Res. 30, 1091–1102.

Davanloo, P., Rosenberg, A.H., Dunn, J.J., Studier, F.W., 1984. Cloning and expression of the gene for bacteriophage T7 RNA polymerase. Proc Natl Acad Sci U S A 81, 2035–2039. 10.1073/pnas.81.7.2035

Eskes, R., Yang, J., Lambowitz, A.M., Perlman, P.S., 1997. Mobility of yeast mitochondrial group II introns: engineering a new site specificity and retrohoming via full reverse splicing. Cell 88, 865–874. 10.1016/s0092-8674(00)81932-7

Ferat, J.-L., Le Gouar, M., Michel, F., 2003. A group II intron has invaded the genus Azotobacter and is inserted within the termination codon of the essential groEL gene. Mol. Microbiol. 49, 1407–1423.

Figueroa-Bossi, N., Balbontín, R., Bossi, L., 2022. Quick Transformation with Plasmid DNA. Cold Spring Harb Protoc 2022, Pdb.prot107854. 10.1101/pdb.prot107854

García-Rodríguez, F.M., Neira, J.L., Marcia, M., Molina-Sánchez, M.D., Toro, N., 2019. A group II intron-encoded protein interacts with the cellular replicative machinery through the β-sliding clamp. Nucleic Acids Res 47, 7605–7617. 10.1093/nar/gkz468

Granlund, M., Michel, F., Norgren, M., 2001. Mutually exclusive distribution of IS1548 and GBSi1, an active group II intron identified in human isolates of group B streptococci. J. Bacteriol. 183, 2560– 2569. 10.1128/JB.183.8.2560-2569.2001

Guo, H., Karberg, M., Long, M., Jones, J.P., Sullenger, B., Lambowitz, A.M., 2000. Group II introns designed to insert into therapeutically relevant DNA target sites in human cells. Science 289, 452–457. 10.1126/science.289.5478.452

Haack, D.B., Yan, X., Zhang, C., Hingey, J., Lyumkis, D., Baker, T.S., Toor, N., 2019. Cryo-EM Structures of a Group II Intron Reverse Splicing into DNA. Cell 178, 612–623.e12. 10.1016/j.cell.2019.06.035

Jacquier, A., Michel, F., 1987. Multiple exon-binding sites in class II self-splicing introns. Cell 50, 17– 29.

Jiménez-Zurdo, J.I., García-Rodríguez, F.M., Barrientos-Durán, A., Toro, N., 2003. DNA target site requirements for homing in vivo of a bacterial group II intron encoding a protein lacking the DNA endonuclease domain. J Mol Biol 326, 413–423. 10.1016/s0022-2836(02)01380-3

Kumar, C.M.S., Mande, S.C., Mahajan, G., 2015. Multiple chaperonins in bacteria--novel functions and non-canonical behaviors. Cell Stress Chaperones 20, 555–574. 10.1007/s12192-015-0598-8

Lambowitz, A.M., Zimmerly, S., 2011. Group II introns: mobile ribozymes that invade DNA. Cold Spring Harb Perspect Biol 3, a003616. 10.1101/cshperspect.a003616

McNeil, B.A., Zimmerly, S., 2014. Novel RNA structural features of an alternatively splicing group II intron from Clostridium tetani. RNA 20, 855–866. 10.1261/rna.042440.113

Meng, Q., Wang, Y., Liu, X.-Q., 2005. An intron-encoded protein assists RNA splicing of multiple similar introns of different bacterial genes. J Biol Chem 280, 35085–35088. 10.1074/jbc.C500328200

Michel, F., Costa, M., Doucet, A.J., Ferat, J.-L., 2007. Specialized lineages of bacterial group II introns. Biochimie 89, 542–553. 10.1016/j.biochi.2007.01.017

Michel, F., Umesono, K., Ozeki, H., 1989. Comparative and functional anatomy of group II catalytic introns--a review. Gene 82, 5–30. 10.1016/0378-1119(89)90026-7

Mohr, G., Ghanem, E., Lambowitz, A.M., 2010. Mechanisms used for genomic proliferation by thermophilic group II introns. PLoS Biol. 8, e1000391. 10.1371/journal.pbio.1000391

Mohr, G., Kang, S.Y.-S., Park, S.K., Qin, Y., Grohman, J., Yao, J., Stamos, J.L., Lambowitz, A.M., 2018. A Highly Proliferative Group IIC Intron from Geobacillus stearothermophilus Reveals New Features of Group II Intron Mobility and Splicing. J Mol Biol 430, 2760–2783. 10.1016/j.jmb.2018.06.019

Mohr, G., Smith, D., Belfort, M., Lambowitz, A.M., 2000. Rules for DNA target-site recognition by a lactococcal group II intron enable retargeting of the intron to specific DNA sequences. Genes Dev. 14, 559–573.

Monachello, D., Lauraine, M., Gillot, S., Michel, F., Costa, M., 2021. A new RNA–DNA interaction required for integration of group II intron retrotransposons into DNA targets. Nucleic Acids Research 49, 12394–12410. 10.1093/nar/gkab1031

Moran, J.V., Zimmerly, S., Eskes, R., Kennell, J.C., Lambowitz, A.M., Butow, R.A., Perlman, P.S., 1995. Mobile group II introns of yeast mitochondrial DNA are novel site-specific retroelements. Mol. Cell. Biol. 15, 2828–2838.

Novikova, O., Smith, D., Hahn, I., Beauregard, A., Belfort, M., 2014. Interaction between conjugative and retrotransposable elements in horizontal gene transfer. PLoS Genet. 10, e1004853. 10.1371/journal.pgen.1004853

Pfreundt, U., Hess, W.R., 2015. Sequential splicing of a group II twintron in the marine cyanobacterium Trichodesmium. Sci Rep 5, 16829. 10.1038/srep16829

Robart, A.R., Seo, W., Zimmerly, S., 2007. Insertion of group II intron retroelements after intrinsic transcriptional terminators. Proc Natl Acad Sci U S A 104, 6620–6625. 10.1073/pnas.0700561104

Roncarati, D., Scarlato, V., 2017. Regulation of heat-shock genes in bacteria: from signal sensing to gene expression output. FEMS Microbiol Rev 41, 549–574. 10.1093/femsre/fux015

Saldanha, R., Chen, B., Wank, H., Matsuura, M., Edwards, J., Lambowitz, A.M., 1999. RNA and protein catalysis in group II intron splicing and mobility reactions using purified components. Biochemistry 38, 9069–9083. 10.1021/bi982799l

Simon, D.M., Clarke, N.A.C., McNeil, B.A., Johnson, I., Pantuso, D., Dai, L., Chai, D., Zimmerly, S., 2008. Group II introns in eubacteria and archaea: ORF-less introns and new varieties. RNA 14, 1704– 1713. 10.1261/rna.1056108

Singh, N.N., Lambowitz, A.M., 2001. Interaction of a group II intron ribonucleoprotein endonuclease with its DNA target site investigated by DNA footprinting and modification interference. J Mol Biol 309, 361–386. 10.1006/jmbi.2001.4658

Toor, N., Hausner, G., Zimmerly, S., 2001. Coevolution of group II intron RNA structures with their intron-encoded reverse transcriptases. RNA 7, 1142–1152.

Waldern, J., Schiraldi, N.J., Belfort, M., Novikova, O., 2020. Bacterial Group II Intron Genomic Neighborhoods Reflect Survival Strategies: Hiding and Hijacking. Mol Biol Evol 37, 1942–1948. 10.1093/molbev/msaa055

Zhuang, F., Karberg, M., Perutka, J., Lambowitz, A.M., 2009a. EcI5, a group IIB intron with high retrohoming frequency: DNA target site recognition and use in gene targeting. RNA 15, 432–449. 10.1261/rna.1378909

Zhuang, F., Mastroianni, M., White, T.B., Lambowitz, A.M., 2009b. Linear group II intron RNAs can retrohome in eukaryotes and may use nonhomologous end-joining for cDNA ligation. Proc. Natl. Acad. Sci. U.S.A. 106, 18189–18194. 10.1073/pnas.0910277106

Zimmerly, S., Guo, H., Eskes, R., Yang, J., Perlman, P.S., Lambowitz, A.M., 1995a. A group II intron RNA is a catalytic component of a DNA endonuclease involved in intron mobility. Cell 83, 529–538.

Zimmerly, S., Guo, H., Perlman, P.S., Lambowitz, A.M., 1995b. Group II intron mobility occurs by target DNA-primed reverse transcription. Cell 82, 545–554.

Zimmerly, S., Hausner, G., Wu Xc., 2001. Phylogenetic relationships among group II intron ORFs. Nucleic Acids Res 29, 1238–1250. 10.1093/nar/29.5.1238

